# Human-Specific Suppression of Hepatic Fatty Acid Catabolism by RNA-Binding Protein HuR

**DOI:** 10.1101/2025.05.22.655551

**Authors:** Shohei Takaoka, Marcos E Jaso-Vera, Xiangbo Ruan

**Author notes:** **Corresponding:** Xiangbo Ruan, Ph.D. Assistant Professor of Medicine The Johns Hopkins University School of Medicine Division of Endocrinology, Diabetes and Metabolism Johns Hopkins All Children’s Hospital, Institute for Fundamental Biomedical Research 600 Fifth Street S. St. Petersburg, FL 33701.

## Abstract

RNA binding proteins (RBPs) play essential roles in all major steps of RNA processing. Genetic studies in human and mouse models support that many RBPs are crucial for maintaining homeostasis in key tissues/organs, but to what extent the function of RBPs is conserved between humans and mice is not clear. Our recent study using a chimeric humanized liver mouse model found that knocking down human HuR in human hepatocytes resulted in a broad upregulation of human genes involved in fatty acid catabolism. This regulation is human-specific, as the knocking down of mouse HuR in the liver of traditional mouse models did not show these effects. To further study this human-specific role of HuR, we co-overexpressed HuR with PPARα, a master transcription factor that promotes fatty acid catabolism, in cultured cells. We found that HuR suppressed the expression of PPARα induced fatty acid catabolism genes in human cells but not in mouse cells. We provide evidence supporting that the human-specific suppressive effect of HuR is independent of PPARα expression or location. The regulatory effects of HuR are also independent of its role in regulating mRNA stability. Using the human HMGCS2 gene as an example, we found that the suppressive effect of HuR cannot be explained by decreased promoter activity. We further provide evidence supporting that HuR suppresses the pre-mRNA processing of HMGCS2 gene, leading to accumulated intron/pre-mRNA expression of HMGCS2 gene. Furthermore, overexpression of HuR blocked and knocking down of HuR sensitized PPARα agonist-induced gene expression. By analyzing published RNA-seq data, we found compromised pre-mRNA processing for fatty acid catabolism genes in patients with fatty liver diseases, which was not observed in mouse fatty liver disease models. Our study supports the model that HuR suppresses the expression of fatty acid catabolism genes by blocking their pre-mRNA processing, which may partially explain the mild effects of PPARα agonists in treating fatty liver diseases in humans as compared with studies in mice.

## INTRODUCTION

RNA-binding proteins play a central role in regulating transcription, pre-mRNA processing, transporting, translation and decay. Many RBPs bind to the non-coding region of pre-mRNA or mature mRNA to function. As these non-coding regions are much less conserved than protein-coding regions, it is speculated that significant RBP-RNA binding events could be species-specific, and thus, RBPs may have human-specific roles. Human antigen R (HuR) is a ubiquitously expressed RNA-binding protein with an established role in regulating RNA stability, pre-mRNA splicing and RNA transporting^1, 2^. Previous studies in mice with genetic depletion of HuR in hepatocytes support that HuR is essential for hepatic metabolic homeostasis^3^. Loss of HuR promotes diet-induced metabolic-associated fatty liver disease (MAFLD) progression, supporting that HuR protects against fatty liver diseases in mice^4–6^. However, our recent work using a chimeric humanized liver mouse model, where human hepatocytes can replace mouse hepatocytes by more than 90%, revealed that depletion of HuR in human hepatocytes resulted in broad upregulation of human genes involved in fatty acid catabolism, which is very different from those observed in traditional mice with hepatic depletion of mouse HuR^7^. Our result suggests a human-specific role of HuR in regulating hepatic gene expression, and HuR may serve as a negative regulator of fatty acid catabolism in human livers^7^.

In this study, we provide mechanism studies underlying the human-specific role of HuR. We first successfully re-captured the effects of HuR in suppressing PPARα induced fatty acid catabolism genes in cultured human cells. More importantly, this suppressive effect was not observed in cultured mouse cells in the same setting. Through a series of experiments, we provide evidence that the human-specific role of HuR is not related to differences between human and mouse PPARα or between human and mouse promoters of fatty acid catabolism genes. Using human HMGCS2 gene as an example, we further excluded that HuR promotes the degradation of mRNA transcripts. Through careful characterization of the pre-mRNA processing upon induction by PPARα agonists, we found that HuR blocked the HMGCS2 pre-mRNA processing at the last few intron-exon junctions. Finally, we show that HuR overexpression totally blocked, while knocking of HuR sensitized PPARα agonist to induce the expression of fatty acid catabolism genes, indicating HuR could be the key factor explaining mild effects of PPARα agonists in treating fatty liver diseases in human populations.

## RESULTS

### Human-specific suppression of fatty acid catabolism gene expression by HuR

We recently reported that knocking down of human HuR in mice with humanized livers resulted in a human gene profile very different from gene expression profile generated in regular mice with hepatic depletion of mouse HuR^7^. To further characterize the role of HuR in human hepatocytes, we re-analyzed the RNA-seq data and performed GO term analysis. We found that HuR knocking down in humanized livers resulted in a robust upregulation of genes involved in lipid and organic acid catabolic process (**Figure 1**), as represented by genes including ACAA2, ACADM, ACOT2, ACSL1, PDK4 and HMGCS2 (**Table 1**). As PPARα is the master regulator of hepatic lipid catabolic process and many of the HuR knocking down induced genes are established PPARα targets^8^, we speculate that HuR may specifically suppress PPARα induced fatty acid catabolism genes in humans. To experimentally test this, we co-overexpressed human HuR and human PPARα in 293A cells and examined the expression of fatty acid oxidation-related genes. The reason that we started our experiments in 293A cells is because of their human origin, high transfection efficiency, and very low expression of endogenous PPARα. We confirmed that overexpression of PPARα induced the expression of PDK4, HMGCS2, and ACADM. We found that this induction was significantly suppressed by co-overexpressed HuR (**Figures 2A-2C**).

**Figure 1.**
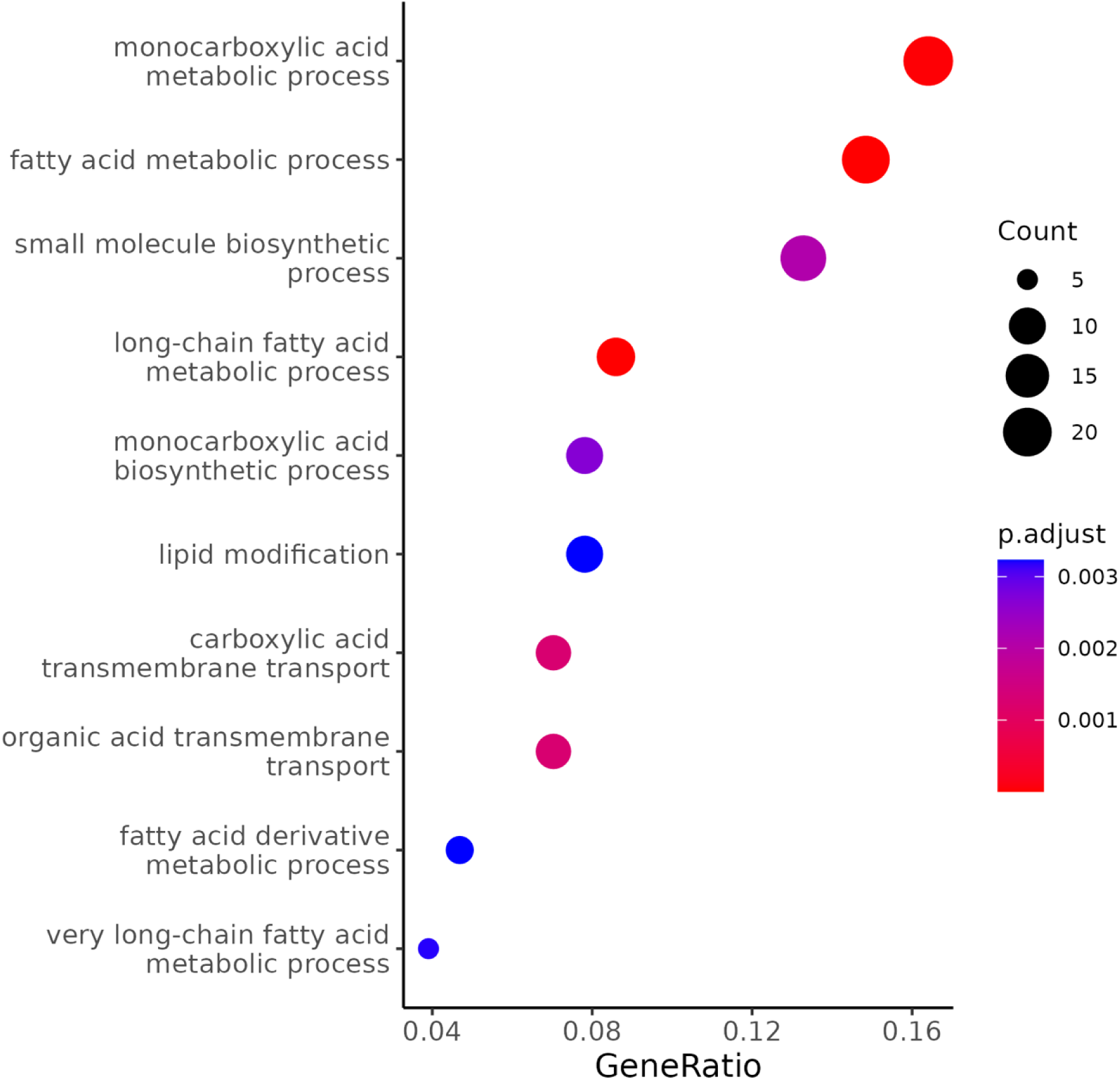
GO term analysis for upregulated genes in humanized livers with knocking down of human HuR.

**Figure 2.**
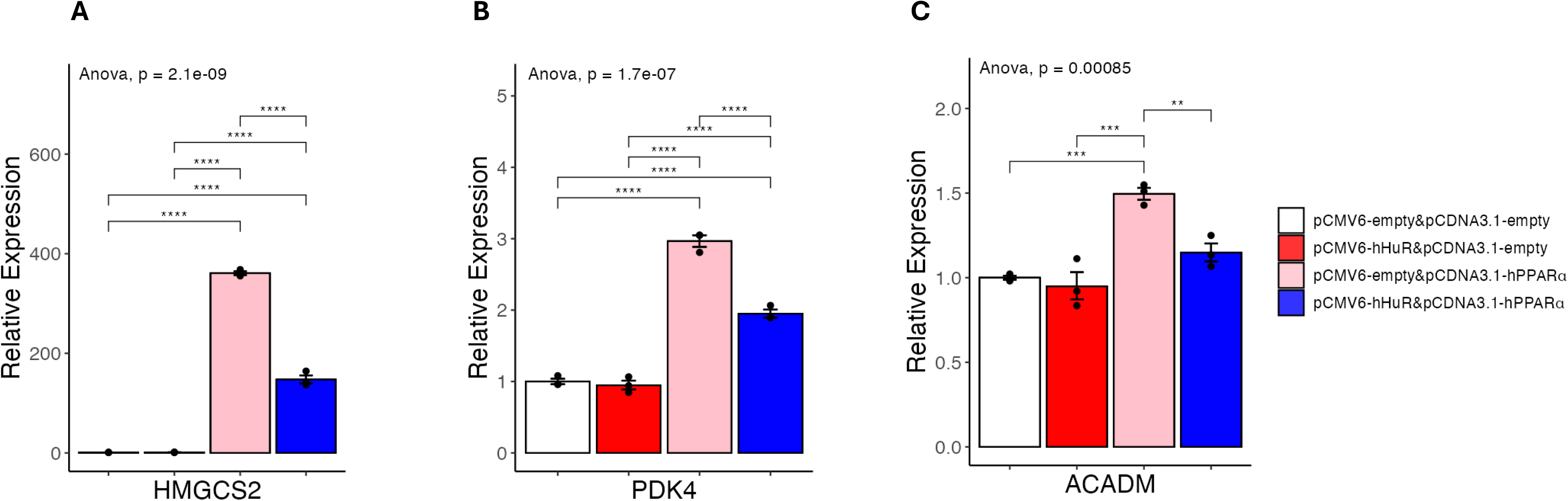
**A-C**, Results of quantitative real-time PCR of mRNAs under hHuR and/or hPPARα overexpression in 293A cells. Data are shown as the geometric mean ± SEM. One-way ANOVA statistical analysis and Tukey HSD test were performed to calculate p-values. *, p<0.05, **, p<0.01, ***, p<0.001, ****, p<0.0001.

**Table 1.** Representative genes involved in fatty acid catabolism that were upregulated by knocking down human HuR in humanized livers.

| Gene_name | Description | Log2FC | p-value |
| --- | --- | --- | --- |
| ACAA2 | acetyl-CoA acyltransferase 2 | 0.607 | 1.13E-05 |
| ACADM | acyl-CoA dehydrogenase medium chain | 0.654 | 1.48E-04 |
| ACOT2 | acyl-CoA thioesterase 2 | 0.681 | 4.64E-04 |
| ACSL1 | acyl-CoA synthetase long chain family member 1 | 0.912 | 1.05E-08 |
| ACSL5 | acyl-CoA synthetase long chain family member 5 | 1.113 | 4.77E-08 |
| HMGCS2 | 3-hydroxy-3-methylglutaryl-CoA synthase 2 | 0.784 | 1.18E-09 |
| PDK4 | pyruvate dehydrogenase kinase 4 | 0.482 | 2.49E-03 |
| SLC27A2 | solute carrier family 27 member 2 | 0.672 | 1.17E-03 |
| SLC27A5 | solute carrier family 27 member 5 | 1.264 | 3.91E-06 |

To test the possibility that HuR may suppress human PPARα but not mouse PPARα-induced gene expression, we performed another co-overexpression experiment in 293A cells to test if HuR can suppress mouse PPARα induced gene expression. As shown in **Figure 3**, mouse PPARα overexpression also resulted in robust upregulation of human HMGCS2 and PDK4 in 293A cells, and these inductions still can be efficiently suppressed by co-overexpressed HuR. This result supports that HuR’s human-specific suppressive effects are not due to differences between human and mouse PPARα.

**Figure 3.**
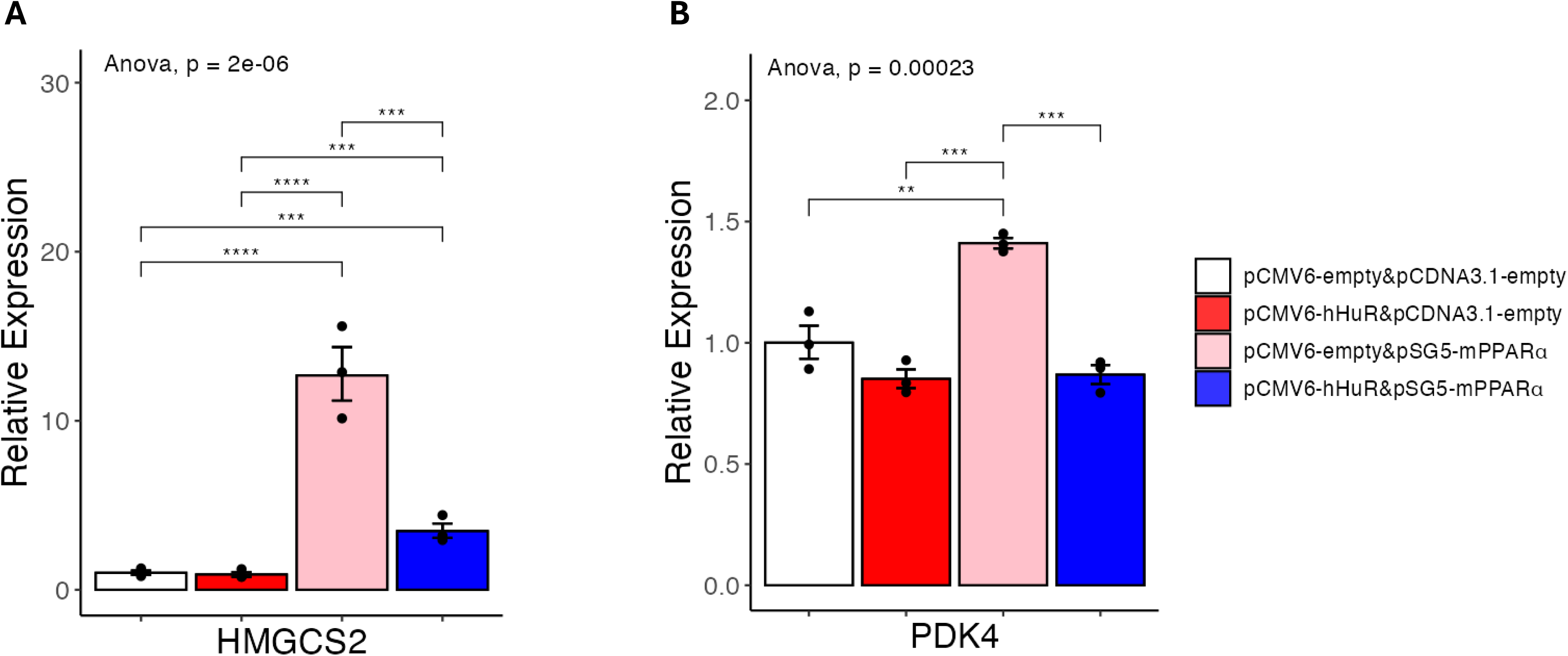
**A-B**, Results of quantitative real-time PCR of mRNAs under hHuR and/or mPPARα overexpression in 293A cells. Data are shown as the geometric mean ± SEM. One-way ANOVA statistical analysis and Tukey HSD test were performed to calculate p-values. *, p<0.05, **, p<0.01, ***, p<0.001, ****, p<0.0001.

To further test if the species-specific role of HuR can be captured in cultured cells, we performed a similar co-overexpression experiment in the mouse NMuMG cells due to its relatively high transfection efficiency. As shown in **Figure 4**, while the PPARα target Hmgcs2 and Pdk4 can be efficiently induced in this setting, co-overexpression of HuR showed no effects in suppressing PPARα-induced mouse Hmgcs2 expression. Together with our in vivo studies, these in vitro data support HuR specifically suppresses the expression of fatty acid catabolism genes in human cells.

**Figure 4.**
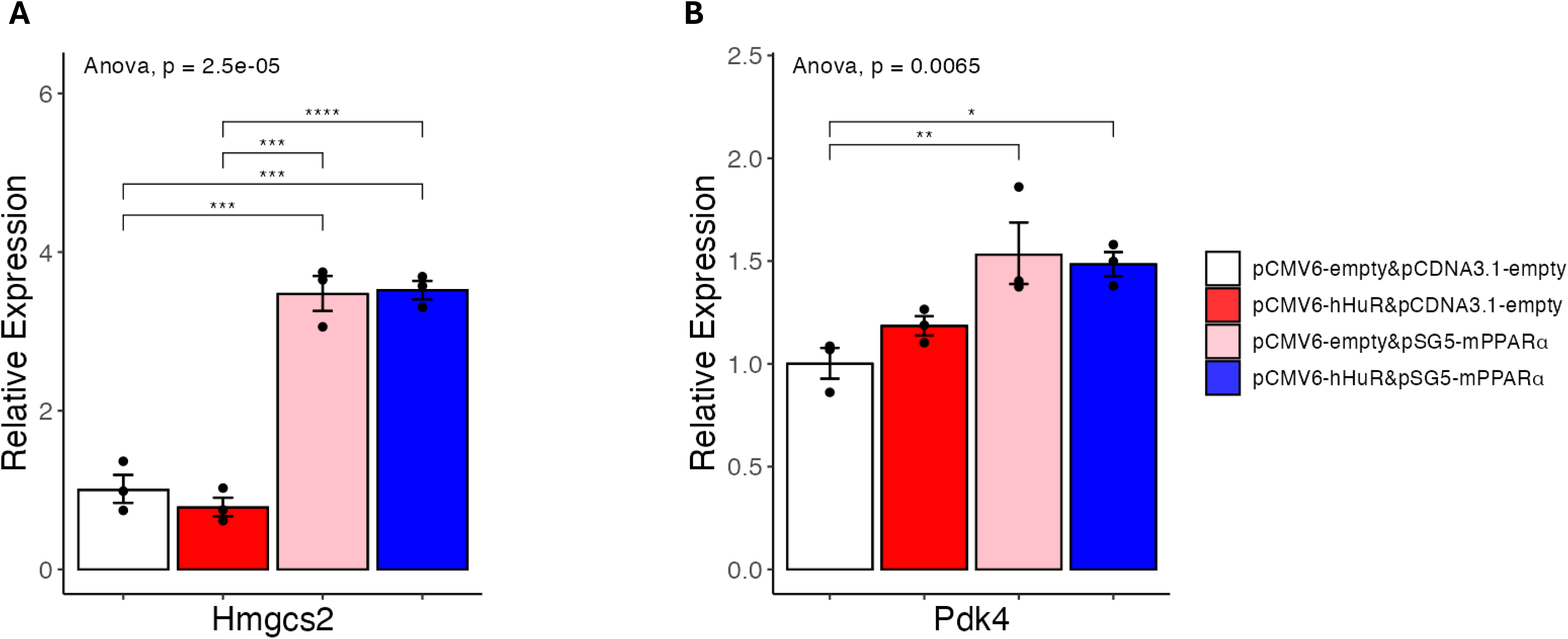
**A-B**, Results of quantitative real-time PCR of mRNAs under hHuR and/or mPPARα overexpression in NMuMG cells. Data are shown as the geometric mean ± SEM. One-way ANOVA statistical analysis and Tukey HSD test were performed to calculate p-values. *, p<0.05, **, p<0.01, ***, p<0.001, ****, p<0.0001.

### HuR suppresses fatty acid catabolism independent of PPARα

To explore the potential mechanism of HuR-mediated suppression of fatty acid catabolism in human cells, we first checked if HuR directly suppresses human PPARα expression. Unexpectedly, as shown in **Figures 5A-5B**, HuR overexpression led to a higher expression of human PPARα at mRNA levels, as compared with PPARα overexpression alone. Western blot analysis confirmed that HuR overexpression led to increased PPARα protein levels (**Figure 5C**). This result suggests that the suppressive role of HuR is independent of PPARα expression. We also checked PPARα expression in our humanized liver RNA-seq and found knocking down of human HuR showed no effects on the mRNA expression of human PPARα (**Figure 5D**), suggesting that increased PPARα mRNA and protein upon HuR overexpression in vitro could be related to the general RNA stabilizing effects of HuR.

**Figure 5.**
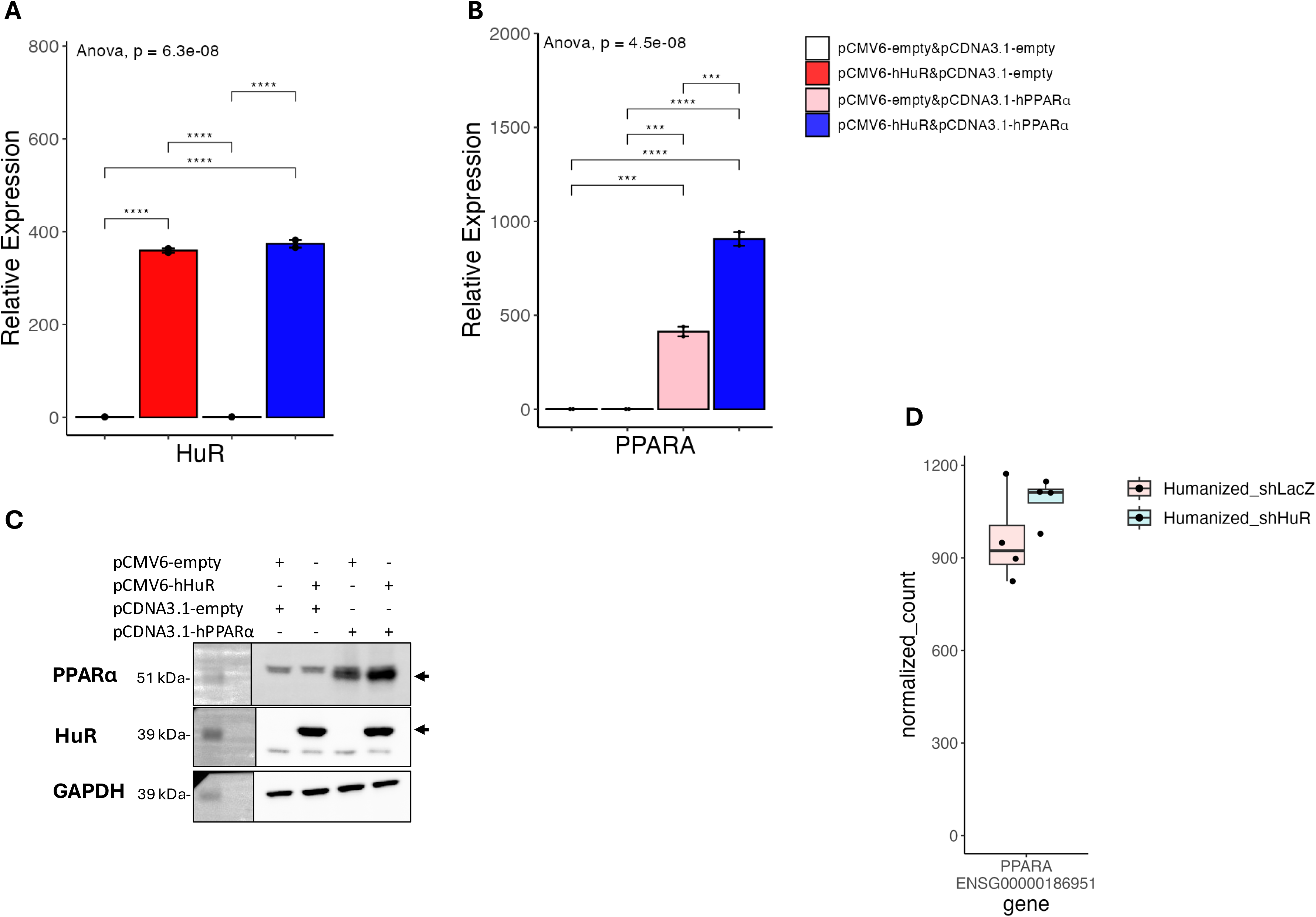
**A-B**, Results of quantitative real-time PCR of mRNAs under hHuR and/or hPPARα overexpression in 293A cells. Data are shown as the geometric mean ± SEM. One-way ANOVA statistical analysis and Tukey HSD test were performed to calculate p-values. **C**, Western blot analysis confirming the overexpression of HuR and PPARα in 293A cells. GAPDH was used as a loading control. Overexpressed HuR and PPARα were pointed out by arrows. Molecular weight markers are indicated on the left. **D**, Normalized count plot showing the expression levels of PPARα in humanized mouse liver samples following HuR knockdown. *, p<0.05, **, p<0.01, ***, p<0.001, ****, p<0.0001.

It was recently reported that hepatic PPARα may undergo nuclear-cytoplasmic shuttling^9^, and HuR is known to translocate between the nucleus and the cytoplasm of a cell^10^. It is possible that HuR overexpression led to increased total PPARα but decreased nuclear PPARα. To test this, we performed nuclear/cytosol fractionation in 293A cells. We found that HuR overexpression resulted in proportionally higher expression of PPARα in both cytosol and nuclear fraction (**Figure 6**). Our results support that the suppressive effect of HuR is not dependent on the expression or location of PPARα.

**Figure 6.**
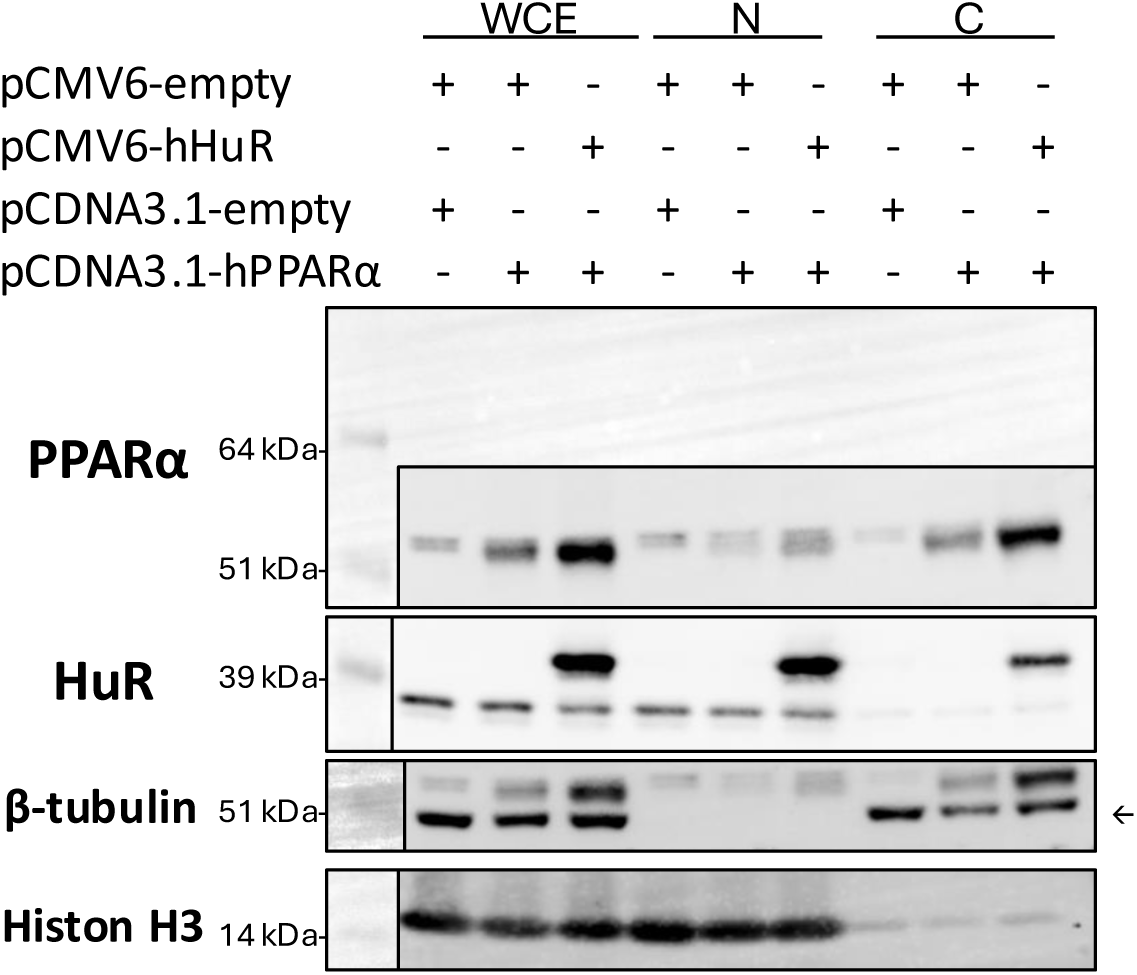
Western blot analysis of nuclear and cytoplasmic fractions from 293A cells overexpressing hHuR and PPARα. Fractionation was performed to assess the subcellular distribution of PPARα in the presence of HuR overexpression. Histon H3 and β-tubulin were used as nuclear and cytoplasmic markers, respectively. β-tubulin was rebloted from the PPARα membrane (the β-tubulin band was pointed out by arrow). WCE, whole cell extract; N, nuclear fraction; C, cytosol fraction.

### Human-specific role of HuR is independent of mRNA stability or promoter activity

Since HuR is an RNA-binding protein with an established role in the regulation of RNA stability^1^, we next checked if HuR expression promotes the degradation of PPARα induced genes. As shown in **Figures 7A-7C**, the decay rates of HMGCS2, PDK4, and ACADM were not significantly different between PPARα overexpression and PPARα plus HuR overexpression groups. This result supports that HuR suppresses PPARα-induced gene expression independent of its role in regulating mRNA stability. These observations, together with the fact that the suppression of PPARα target genes by HuR is human-specific, make us speculate that HuR acts through evolutionarily poorly conserved factors, such as human-unique DNA sequences at the promoter region, to suppress PPARα induced gene transcription. To test this, we cloned a 2 kb promoter for both human and mouse HMGCS2 gene and performed a luciferase reporter assay (**Figure 8A**). As shown in **Figures 8B-8C**, we found that, as previously reported, both mouse and human HMGCS2 promoters can be robustly induced by PPARα. HuR overexpression, however, only resulted in a less than 10 percent suppression of PPARα induced-human HMGCS2 promoter activity (**Figure 8B**). In contrast, HuR overexpression led to a 27% suppression of PPARα induced mouse HMGCS2 promoter activity (**Figure 8C**). We noticed that mouse HMGCS2 promoter showed a higher activity as compared with human HMGCS2 promoter upon PPARα expression, in line with the prediction that mouse HMGCS2 promoter has two PPARα/RXRα binding sites and human HMGCS2 promoter only has one. Indeed, the luciferase reporter^11^ that contains three artificial Peroxisome Proliferator Response Element (PPRE) was highly induced by PPARα, and HuR overexpression led to a 28% suppression of this reporter (**Figure 8D**). Taken together, our reporter assay suggests that while HuR possesses a weak suppressive effect on PPRE-mediated transcription activity, this effect is unlikely to explain the human-specific, and strong suppression of endogenous human HMGCS2 expression by HuR.

**Figure 7.**
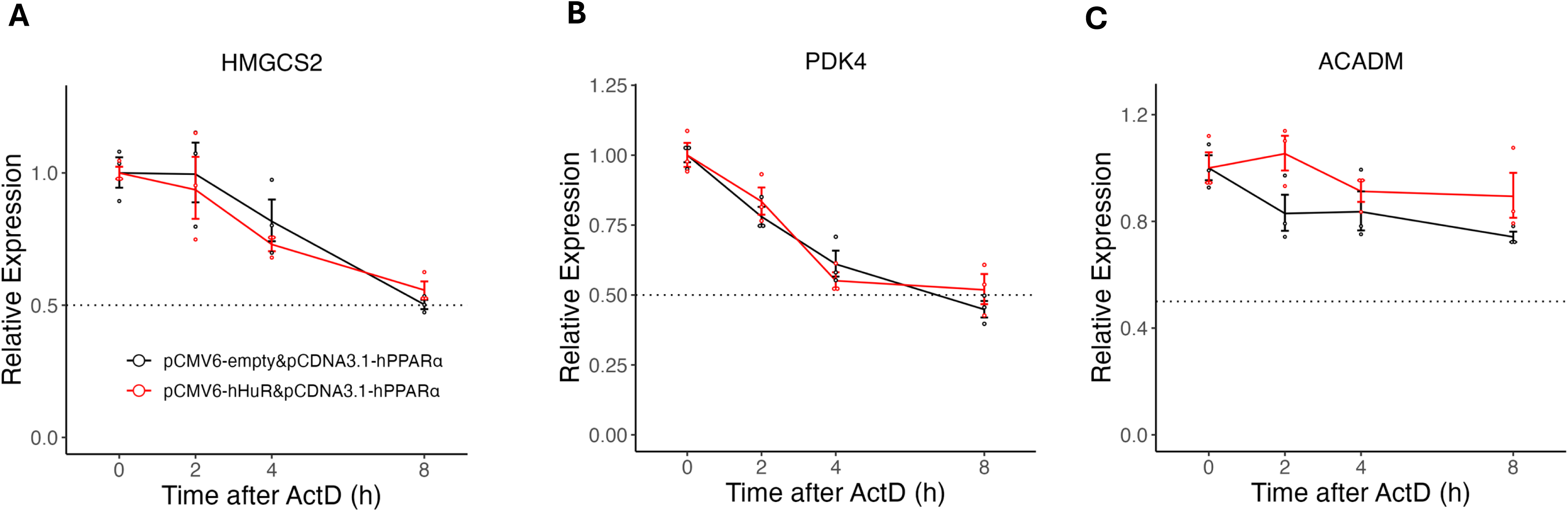
**A-C**, qPCR analysis of mRNA decay kinetics following Actinomycin D (ActD) treatment in 293A cells overexpressing hPPARα alone or co-overexpressing hHuR and hPPARα. Total RNA was extracted at indicated time points (0, 2h, 4h, and 8h) after ActD treatment, and relative mRNA levels of target genes were quantified using qPCR. Gene expression was normalized to β-actin signal, and data were further adjusted so that the relative expression at time 0 was set to 1. Data are shown as the geometric mean ± SEM. One-way ANOVA statistical analysis and Tukey HSD test were performed to calculate p-values.

**Figure 8.**
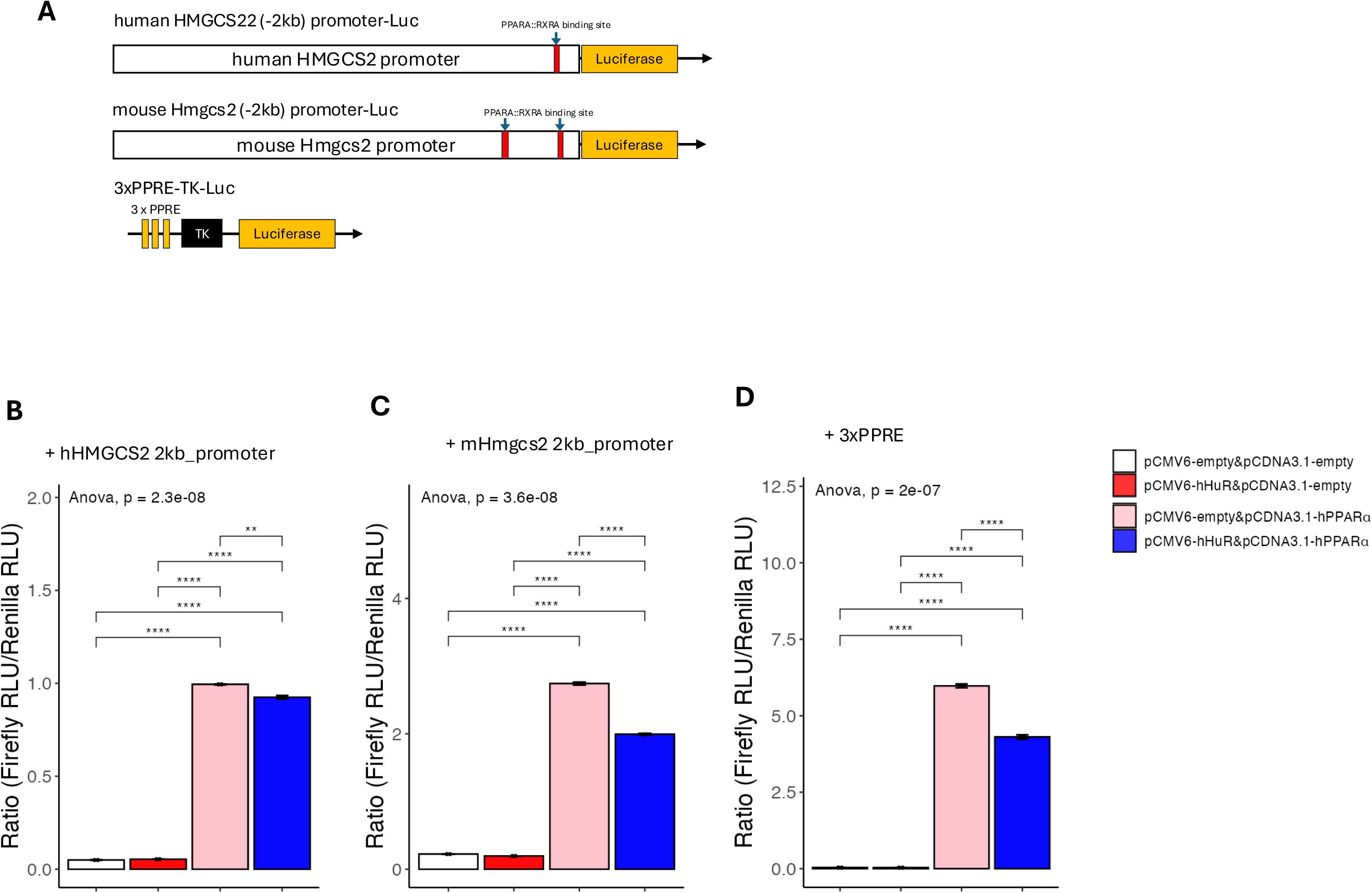
**A**. Diagram depicting the core components of the luciferase plasmid cloned from the 2 kb upstream promoter region of human and mouse HMGCS2. Predicted PPARA-RXRA binding sites were identified using the JASPAR database and are indicated on the schematic. **B-D**, Luciferase reporter assay examining the effects of hHuR and hPPARα on the HMGCS2 promoter and a synthetic PPAR response element (PPRE). Data are shown as the mean ± SEM. One-way ANOVA statistical analysis and Tukey HSD test were performed to calculate p-values. *, p<0.05, **, p<0.01, ***, p<0.001, ****, p<0.0001.

### HuR blocks pre-mRNA processing of PPARα target genes

Given the observation that the human-specific suppressive role of HuR could not be explained by the promoter activity of its target genes, we next asked if HuR regulates the alternative pre-mRNA splicing of the fatty acid catabolism genes. We first checked our RNA-seq data in the humanized liver mouse model and found no evidence of alternative splicing of HMGCS2, PDK4, or ACADM (**Supplementary Figure 1**). However, given that pre-mRNA processing is highly coupled to gene transcription^12, 13^, we next aimed to further determine if HuR affects the pre-mRNA processing using the HMGCS2 gene as an example. We thus designed an array of real-time PCR primers amplifying introns or intron-exon junctions of human HMGCS2 genes covering the whole gene body (**Figure 9A**). We then checked the expression of the regions amplified by these primers as in the setting of **Figure 2**. As expected, compared with control, PPARα overexpression alone significantly induced the expression of all the HMGCS2 transcript regions amplified by our tilling primers, supporting a general transcriptional activation (**Figures 9B-9H**). However, we found that while the expression of primers amplifying the first few introns/intron-exon junctions was suppressed by HuR overexpression, the primers amplifying the last few introns/intron-exon junctions were not. Indeed, the suppressive effects of HuR at the pre-mRNA level gradually decreased from the transcription start site to the transcription termination site (**Figures 9B-9H**). Furthermore, regions amplified by primers covering intron9 and intron8-exon9 junction showed a trend of increased expression by HuR overexpression alone (**Figures 9G-9H**). These data indicate that HuR suppresses the 3’ end pre-mRNA processing of HMGCS2, leading to accumulated pre-mRNA not being efficiently processed to mature mRNA.

**Figure 9.**
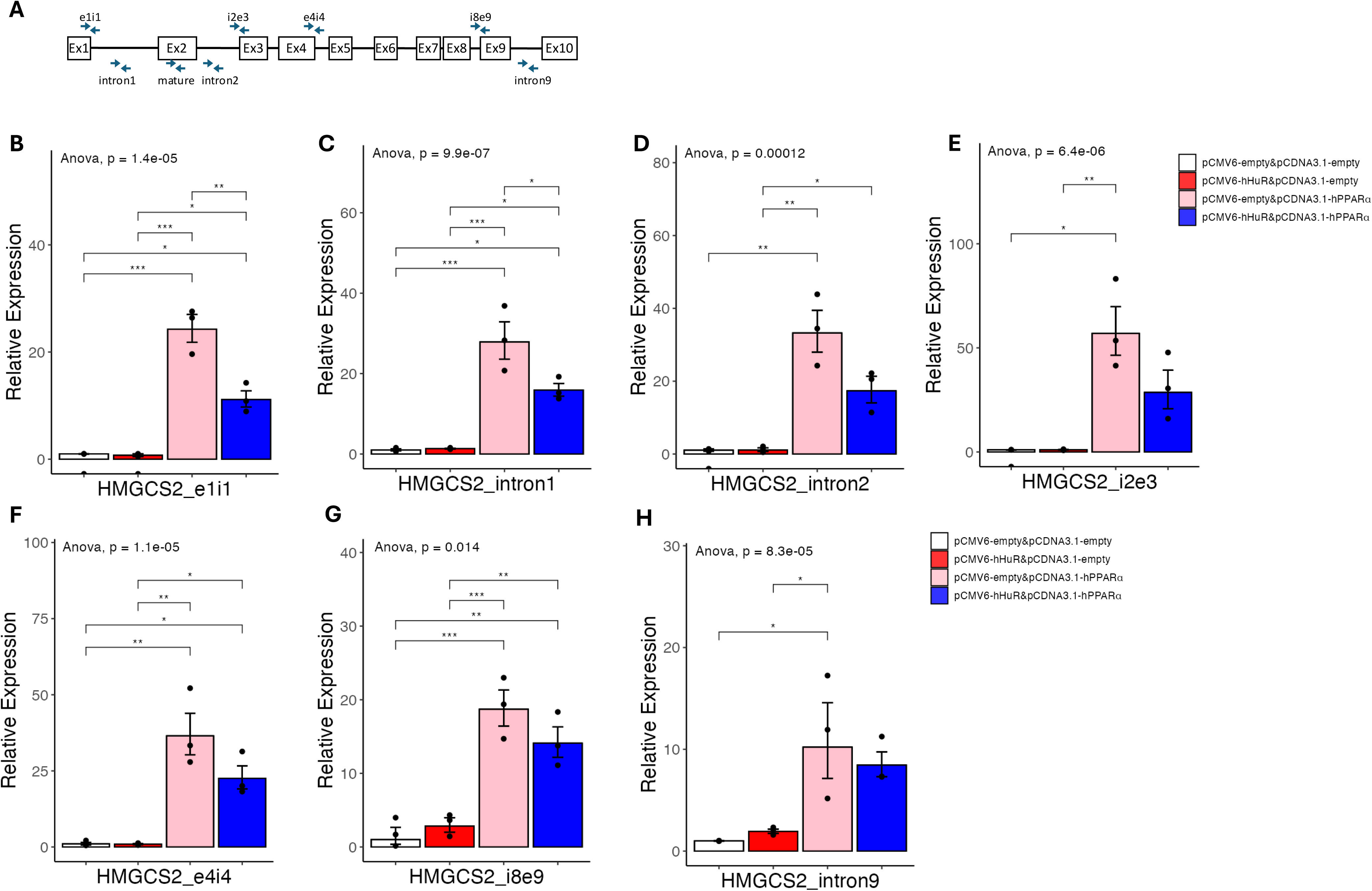
**A**. Diagram showing human HMGCS2 gene structure and the design of pre-mRNA primers. The primer detecting mature mRNA of HMGCS2 was also marked. **B-H**. Results of quantitative real-time PCR of HMGCS2 pre-mRNAs under hHuR and/or hPPARα overexpression in 293A cells. Data are shown as the geometric ± SEM. One-way ANOVA statistical analysis and Tukey HSD test were performed to calculate p-values. *, p<0.05, **, p<0.01, ***, p<0.001.

### HuR suppresses PPARα agonist-induced gene expression

To further study the pre-mRNA processing of HMGCS2, we next treated 293A cells with a specific PPARα agonist, GW7647. We reason that ligand-dependent activation of PPARα allows us to better capture the dynamics of HMGCS2 pre-mRNA processing. To this end, we co-transfected HuR and mouse PPARα in 293A cells and titrated the cells with GW7647 for 5 hours. We used mouse PPARα in this setting due to relatively lower activity as compared to human PPARα (**Figure 2 and Figure 3**), thus allow us to have a large window to capture the effects of GW7647. As shown in **Figure 10A**, GW7647 does dependently induced the expression of HMGCS2 mature mRNA levels in cells with overexpression of mPPARα alone, and co-overexpressed HuR significantly suppressed HMGCS2 expression at all doses. At the pre-mRNA level, although with variations among the technical repeats, the expression patterns of pre-mRNA regions covered by exon1-intron1, intron2, and intron2-exon3 primers were similar to mature mRNA, while the expression patterns of pre-mRNA regions covered by intron8-exon9 and intron9 were different (**Figures 10B-10G**). Specifically, pre-mRNA regions covered by intron8-exon9 and intron9 showed, on average, increased expression at the basal level and cannot be further induced by GW7647 (**Figures 10F-10G)**. These results further support that HuR-mediated suppression of pre-mRNA processing, rather than gene transcription, is the upstream event leading to the downregulation of PPARα agonist-induced gene expression.

**Figure 10.**
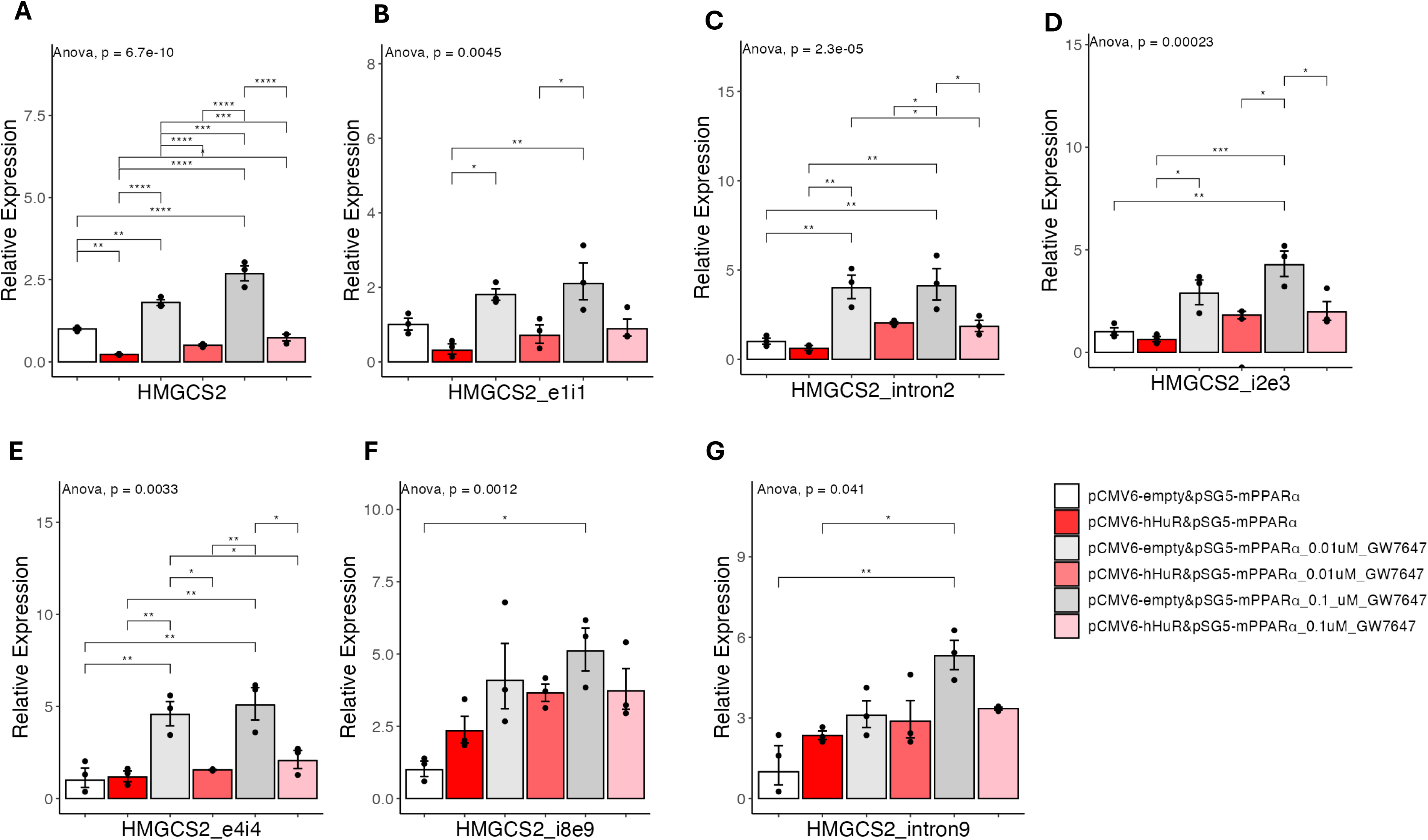
**A-G**. Results of quantitative real-time PCR of HMGCS2 pre-mRNAs under hHuR and/or mPPARα overexpression treated with GW7647 in 293A cells. Data are shown as the geometric ± SEM. One-way ANOVA statistical analysis and Tukey HSD test were performed to calculate p-values. *, p<0.05, **, p<0.01, ***, p<0.001, ****, p<0.0001.

Our observation raised the possibility that knocking down HuR may sensitize PPARα agonist inducing downstream target genes. To test this, we knocked down HuR in human HepG2 cells, a human liver cancer cell line where HuR is highly expressed^14^, and then tested the does responses of GW7647 (**Figure 11A**). As shown in **Figure 11B**, 0.1 uM GW7647, on average induced more HMGCS2 mature mRNA expression in cells with knocking down of HuR as compared with control. At the pre-mRNA level, regions amplified by primers covering exon1-intron1 and exon4-intron4 showed, on average, increased expression in HuR knocking down group (**Figures 11C-11D**). Interestingly, in the HuR knocking down group, the region amplified by primer covering intron8-exon9 showed, on average, lower expression at basal condition, which was restored upon GW7647 treatment as compared with control (**Figure 11E**). These results are consistent with our overexpression experiments, supporting knocking down of HuR could sensitize cells to PPARα agonist induced HMGCS2 expression.

**Figure 11.**
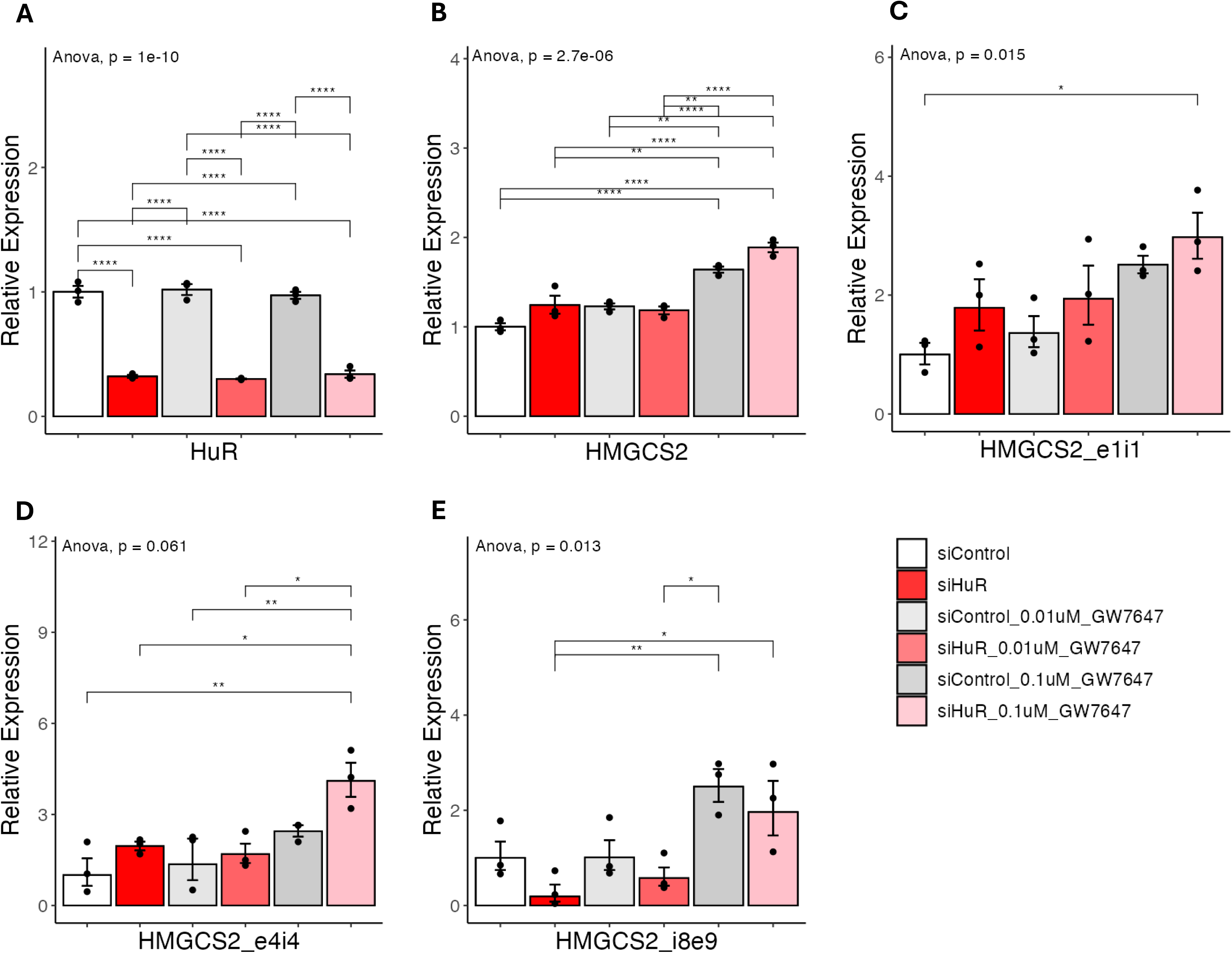
**A-E**. Results of quantitative real-time PCR of HMGCS2 pre-mRNAs under si-hHuR or si-Control in HepG2 cells. Data are shown as the geometric ± SEM. One-way ANOVA statistical analysis and Tukey HSD test were performed to calculate p-values. *, p<0.05, **, p<0.01, ***, p<0.001, ****, p<0.0001.

### Accumulation of HMGCS2/PDK4 pre-mRNAs in human metabolic dysfunction-associated steatohepatitis

To further understand the pathophysiological significance of our finding, we next asked if the suppressed pre-mRNA processing of fatty acid catabolism genes could be captured in human liver samples. To this end, we applied the intron-exon combined RNA-seq analysis pipeline to a published human liver RNA-seq dataset representing gene expression from both healthy individuals and MASH patients^15^. Indeed, HuR was reported to be upregulated in the liver of human MAFLD/MASH patients^3^ but not in mouse MAFLD/MASH^5, 6^. As shown in **Figure 12A**, in line with previous reports showing decreased expression of HMGCS2 in human MASH patients^16^, we found the number of exon-mapped reads was much lower for HMGCS2. However, there were increased intron-mapped reads specifically in the regions from intron 6 to intron 9 regions, but not at the first few introns, in MASH patients as compared with health controls (**Figure 12A)**. A similar accumulation of intron 2/7/10 mapped reads was also observed for PDK4, while its exon-mapped reads were downregulated in MASH (**Figure 12B**). This result suggests compromised pre-mRNA processing leading to pre-mRNA accumulation for fatty acid catabolism genes in human MASH. To verify the compromised pre-mRNA processing for fatty acid catabolism genes is a human-specific regulatory mechanism, we next performed a similar analysis using a liver RNA-seq dataset from mice fed with a high-fat diet (HFD) or normal chow diet for 26 weeks^17^. As shown in **Figures 12C & 12D**, 26 weeks of HFD feeding resulted in decreased both intron and exon mapped reads for Hmgcs2 and Pdk4, supporting downregulated transcription is the major mechanism underlying decreased fatty acid catabolism genes in mouse fatty liver diseases.

**Figure 12.**
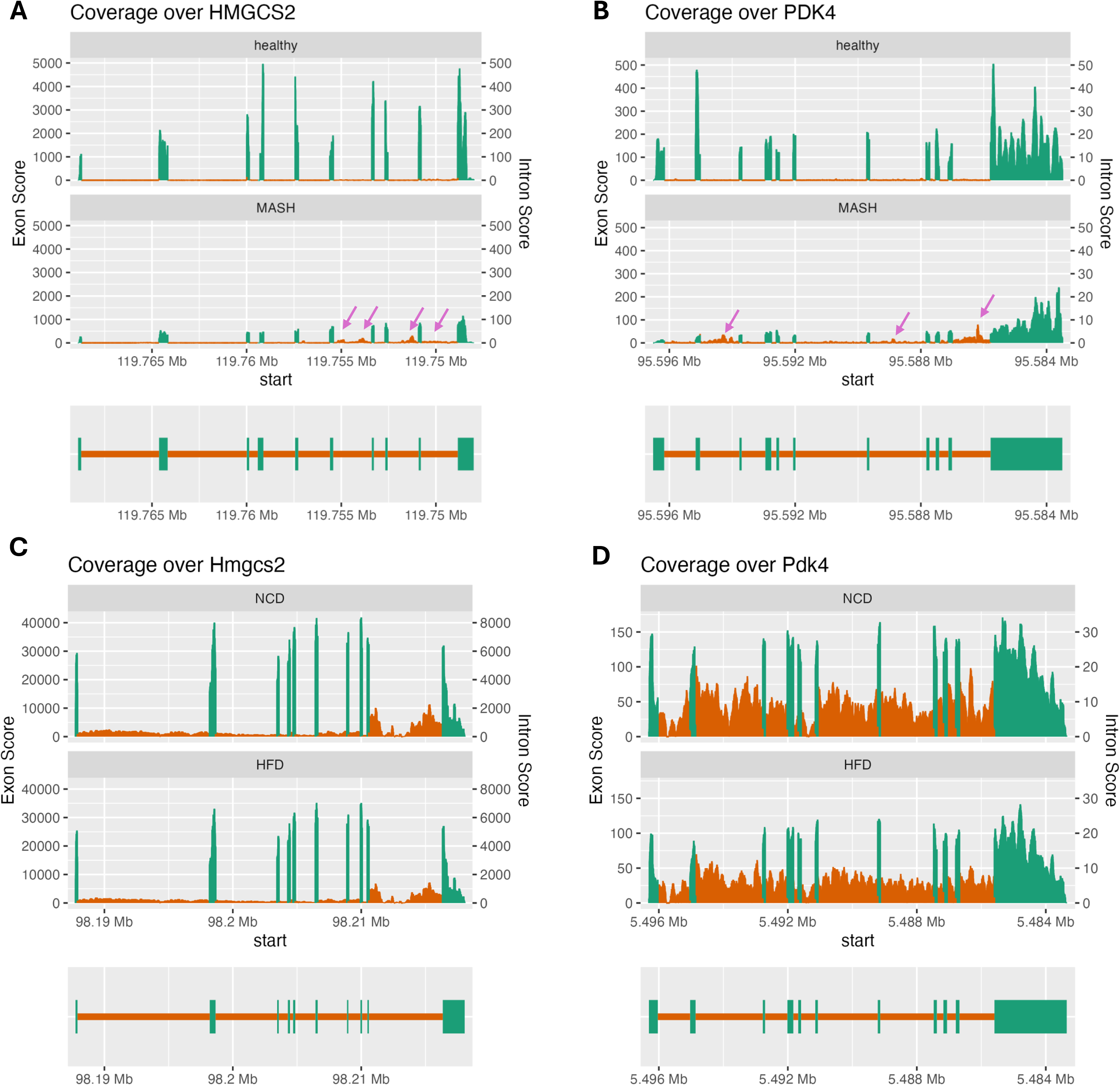
Coverage plots of the indicated genes in human and mouse samples were generated using SuperIntronic. The left Y-axis represents the normalized exon coverage score, while the right Y-axis indicates the normalized intron coverage score. The green regions correspond to exons, and the orange regions represent introns. A gene structure diagram is included at the bottom of each panel, indicating the positions of exonic and intronic regions within each gene and providing a reference for the corresponding RNA-seq read coverage. **A-B**, Human liver RNA-seq data from MASH patients and healthy donors, showing the differential exon and intron coverage for HMGCS2 and PDK4 between the two groups. Regions with accumulated intron-mapped reads were highlighted by purple arrows. **C-D**, Mouse liver RNA-seq data from High-Fat Diet (HFD) fed mice and Normal Chow Diet (NCD) fed control mice, highlighting the differences in exon and intron coverage of Hmgcs2 and Pdk4 in response to diet-induced changes.

## DISCUSSION

In this manuscript, we provide data support that: (1) The function of HuR in human hepatocytes is different from mouse hepatocytes; (2) HuR suppresses the expression of genes involved in fatty acid catabolism, as represented by HMGCS2 and PDK4, likely by blocking the pre-mRNA processing in humans; (3) Compromised pre-mRNA processing contributes to downregulation of fatty acid catabolism genes in human MASH. These findings could potentially explain several previous observations that support human-mouse differences regarding HuR and PPARα biology^8, 18^.

First, recent studies using traditional mouse genetic models support the idea that HuR protects mice from HFD-induced fatty liver diseases. It is interesting that the mechanism of action of HuR varies in these studies, suggesting the complexity of HuR biology^3–6^. We also noticed that among these studies, little information was provided to explore the role of HuR in human livers. Indeed, while HuR was reported downregulated by HFD in mouse livers, studies using human samples support that HuR is upregulated in human MAFLD/MASH^3^, indicating a different role of HuR in the liver between humans and mice. Our data generated using the humanized liver mouse model further support the human-specific role of HuR in suppressing the expression genes involved in fatty acid catabolism, which was not reported in any of the traditional mouse model-based HuR studies^4–6^. Our study thus also supports the significance of utilizing humanized liver mouse models to understand human hepatocyte biology in vivo.

Second, most of the HuR studies largely focused on the role of HuR in regulating mRNA stability. Our mechanism study supports that the regulatory role of HuR on fatty acid catabolism is not related to mRNA stability; instead, HuR blocks the pre-mRNA processing of fatty acid catabolism genes. This is in line with the fact that HuR is highly enriched in the nuclear of human cells. Although gene expression changes at pre-mRNA levels are usually linked to gene transcription, our data support that blocking pre-mRNA processing is the main mechanism underlying HuR’s suppressive effects. First, our luciferase reporter assay demonstrated that HuR has little effect in suppressing human HMGCS2 promoter activity. Second, the expression of regions amplified by primers covering HMGSCS2 gene body showed a different pattern. Specifically, the expression patterns of regions close to the transcription start site were similar to mature mRNA, but the expression patterns of regions close to the last few intron/exons were different from mature mRNA. Pre-mRNA processing and gene transcription are highly coupled processes, where the defect in one process affects the other^12, 13^. We argue that if suppressed gene transcription is the primary mechanism underlying HuR action, we would expect a uniform expression pattern of regions amplified by our tilling primers. As such, together with our luciferase reporter assay, the intron-exon/intron expression analyses support that HuR-mediated blocking of pre-mRNA processing is the upstream event leading to compromised gene transcription for fatty acid catabolism genes in humans.

Third. HuR may explain the limited capacity of PPARα ligand inducing fatty acid catabolism in human livers^19, 20^. As a master regulator of fatty acid catabolism in the liver, PPARα agonist treatment protects mice from diet-induced fatty liver diseases^21–23^. However, several human studies found that the clinically used PPARα agonists have very limited effects in improving MAFLD-associated steatosis^19^. Indeed, fenofibrate, the most widely clinically used PPARα ligand, is for treating hyper-triglyceride, which is mainly mediated by the role of PPARα to induced lipoprotein lipase in the muscle. These observations suggest additional regulatory mechanisms controlling PPARα induced gene expression in human livers. This is in line with previous studies performing PPARα agonist treatment in humanized livers, where PPARa induced a more robust upregulation of fatty acid catabolism genes in mouse cells as compared with those in human hepatocytes^18^. Especially, our RNA-seq analysis further supports our conclusion that suppressed pre-mRNA processing is an important regulatory mechanism contributing to decreased fatty acid catabolism gene expression in humans but not mice (**Figure 12**).

In summary, using data generated in humanized liver mouse models, in vitro cultured cells and human livers, we provide HuR-mediated suppression of pre-mRNA processing as a human-specific regulatory mechanism in suppressing fatty acid catabolism in the liver. Future studies such as Clip-seq^24^ will help to define the pre-mRNA regions/motifs that interact with HuR. More in vivo studies in humanized liver mouse models will allow us to test if blocking HuR could sensitize PPARα ligands to treat fatty liver diseases in humans.

## METHODS

### Cell culture

293A cells (ThermoFisher Cat# R70507) and HepG2 cells (ATCC Cat# HB-0865) were maintained in Dulbecco’s modified Eagle’s medium (DMEM; ThermoFisher Cat# 11965118) supplemented with 10% FBS (ThermoFisher Cat# 26140079) and NMuMG cells (ATCC Cat# CRL1636) were maintained in DMEM supplemented with 10% FBS and 10 µg/ml insulin (Sigma-Aldrich Cat# I0516) at 37 °C in a 5% CO2 humidified incubator. 293A or NMuMG cells were plated in 24-well plates 1 day before plasmid transfection so that confluency was approximately 70% at the time of transfection. Cells were transfected with 400 ng of pCMV6-hHuR (OriGene Cat# RC201562), 100 ng of pCDNA3.1-hPPARα (Addgene Cat# 169019), 100 ng of pSG5-mPPARα (Addgene Cat# 22751), or equal amounts of the corresponding empty plasmid using Lipofectamine 2000 according to the manufacturer’s protocol. The combination of each plasmid is shown in the figure legends. Plasmid-transfected cells were harvested 48 hours after transfection. HepG2 cells were plated in a 24-well plate one day before siRNA transfection so that confluency was approximately 70% at the time of transfection. HepG2 cells were transfected with 50 nM of Lincode Non-targeting Control Pool (Dharmacon Cat# D-001320-10-05) or siRNA targeting HuR (ON-TARGETplus siRNA; Dharmacon Cat# L-003773-00-0005) by using Lipofectamine RNAiMax (ThermoFisher Cat# 13,778–150), following the manufacturer’s instructions. siRNA-transfected cells were harvested 72 hours after transfection.

### Actinomycin D treatment

To measure mRNA decay kinetics, the medium of 293A cells was replaced with complete medium containing 0.5 μg/mL actinomycin D (Millipore Cat# A9415-10MG) 2 days after plasmid transfection. Samples were collected at 0, 2, 4, and 8 hours after treatment.

### GW7647 treatment

In the GW7647 (Sigma-Aldrich, G6793) treatment experiments, the medium was replaced 5 hours before harvest with a complete medium supplemented with different concentrations of GW7647 (untreated, 0.01 µM, 0.1 µM).

### RNA extraction and qPCR

Total RNA was extracted and purified using the Qiagen RNeasy Mini Kit (Qiagen Cat# 74,106) following the manufacturer’s instructions and performing the on-column DNA digestion. 500 ng of total RNA was used for cDNA synthesis with the SuperScript III First-Strand Synthesis SuperMix for qRT-PCR (Invitrogen Cat# 11,752,050). cDNA was diluted 10-fold with RNase-free water. qPCR reactions consisted of 7.5 µl of the Power SYBR Green PCR Master Mix (ThermoFisher Cat# 4,368,702), 1.5 μL of cDNA, and 6 µl of 0.5 µM primers (primers are listed in Table below). qPCR was performed using a ThermoFisher Quantstudio 7 Flex using a 386-well plate (ThermoFisher Cat# 4,309,849) with Standard Mode. Target gene expression was normalized by the ΔΔCT method, and the GAPDH or β-Actin signal was used as an internal control.

### Primer list

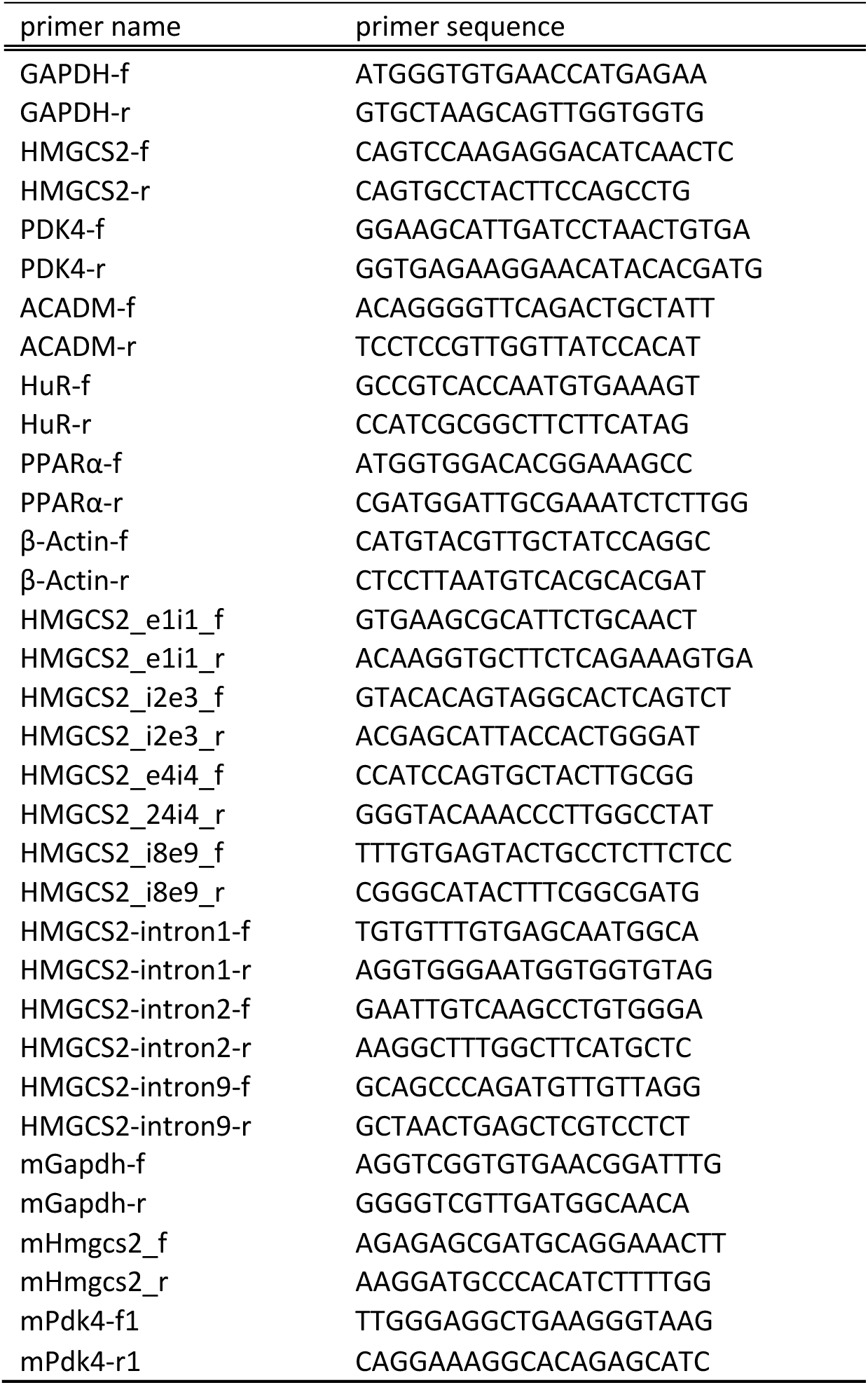

#### Protein extraction, Cell fractionation, and Western blotting

Total protein was extracted using 1xLDS loading buffer (1x NuPAGE LDS Sample Buffer (Invitrogen Cat# NP0007), 5% β-mercaptoethanol (BIO-RAD Cat# 1610710), 1 mM PMSF (Sigma-Aldrich Cat# 93482), Halt™ Protease and Phosphatase Inhibitor Cocktail (ThermoFisher Cat# 78444)). Cell fractionation was performed following the REAP method^25^. Protein lysates were boiled at 70°C for 10 minutes and stored at −20 °C till used. Samples were pre-heated at 37 °C for 5 minutes and loaded to 4–12% NuPAGE™ Bis-Tris Mini Protein

Gels (Invitrogen Cat# NP0322PK2) set in XCell SureLock Mini-Cell (Invitrogen Cat# EI0001) filled with 1x NuPAGE MOPS SDS Running Buffer (Invitrogen Cat# NP000102) and the electrophoresis was done using PowerPac Basic Power Supply (Bio-Rad Cat# 1645050). Proteins in the gel were transferred onto 0.45μm Immobilon-FL transfer membranes (Millipore Cat# IPFL00010) using an XCell II Blot Module (Invitrogen Cat# EI9051) filled with 1x NuPAGE Transfer Buffer (Invitrogen Cat# NP00061) with 0.01% SDS and 5% methanol. After membranes were blocked with Intercept (PBS) Blocking Buffer (LI-COR Cat# 927-70001), proteins of interest were probed with specific primary antibodies and then appropriate Fluorescent Dye-conjugated secondary antibodies diluted in Intercept T20 (TBS) Antibody Diluent (LI-COR Cat# 927-75001). Fluorescent signals were detected using a ChemiDoc MP Imaging System (Bio-Rad Cat# 12003154). The following primary antibodies were used for immunoblotting: PPARα (SantaCruz Cat# sc-398394), HuR (SantaCruz Cat# sc-5261), GAPDH (Cell Signaling Cat# 5174), β-tubulin (Cell Signaling Cat# 2128), and Histon H3 (Cell Signaling Cat# 4499). The following secondary antibodies were used: IRDye 800CW Goat anti-Rabbit IgG Secondary Antibody (LI-COR Cat# 926-32211) and IRDye 800CW Goat anti-Mouse IgG Secondary Antibody (LI-COR Cat# 926-32210).

#### Luciferase Reporter Assay

293A cells were plated in a 24-well plate one day before plasmid transfection to reach ∼ 70% confluency for overexpression. Cells were transfected with 100 ng of pGL3-human-HMGCS2-2kb promoter, pGL3-mouse-Hmgcs2-2kb promoter, or 3xPPRE-TK-Luc (Addgene Cat# 1015), and 100 ng of pRL-TK, 80 ng of pCMV6-hHuR, 20 ng of pCDNA3.1-hPPARα, or equal amounts of the corresponding empty plasmid using Lipofectamine 2000 according to the manufacturer’s protocol. The combination of each plasmid is shown in the figure legends. Cells were harvested 24 hours after transfection. The assay was performed using the Dual-Luciferase® Reporter Assay (Promega Cat# E1910) according to the manufacturer’s protocol.

#### Bioinformatics & bulk RNA-seq analysis

Raw FASTQ files from multiple RNA-seq datasets were analyzed. For human liver transcriptome analysis, data from healthy (n = 14) and MASH (n = 12) patients were obtained from BioProject No. PRJNA523510. For mouse liver transcriptome analysis, six FASTQ files were downloaded from dataset GSE121340, which includes RNA-seq data from 36-week-old control mice (n = 3) fed a chow diet and 36-week-old obese mice (n = 3) fed a high-fat diet (HFD; D12492, Research Diet, 60% fat by calories). Additionally, humanized mouse liver RNA-seq data from shLacz-treated (n = 4) and shHuR-treated (n = 4) samples were obtained from dataset GSE161462.

The Fastq files were downloaded and prepared using the SRA Toolkit (version 2.11.1) [https://www.ncbi.nlm.nih.gov/books/NBK569238/]. Briefly, the prefetch command was used to download the SRA files of the dataset and convert them into fastq format using the fasterq-dump command. The quality of the fastq files was evaluated using FastQC (version 0.11.8) [http://www.bioinformatics.babraham.ac.uk/projects/fastqc/], and adapters and low-quality reads were trimmed or removed using Trimmomatic (version 0.39)^26^. Filtered reads were mapped to the GRCh38 Ensembl human genome for human data and the GRCm39 Ensembl mouse genome for mouse data using STAR (version 2.7.8a)^27^. For the humanized liver mouse dataset, filtered reads were mapped to a merged GRCh38 and GRCm39 genome set. Using SAMtools, we filtered BAM files by extracting reads uniquely mapped to the human genome for subsequent differential gene expression (DEG) analysis. DEG analysis was performed using DESeq2 (version 1.36.0)^28^ to calculate Log2 fold change, p-values using the Wald test. Differentially expressed genes were defined as genes with false discovery rate (FDR) <= 0.05 and |Log2FC| >=0.58. DEGs with top 10,000 in baseMean calculated by DESeq2 were used as an input for gene ontology (GO) enrichment analysis using clusterProfiler (version 4.0.5)^29^. GO terms with FDR <= 0.01 were considered significant. The Normalized counts of PPARα from the humanized liver dataset were extracted using the plotCounts function from DESeq2 for drawing the normalized count plots. Coverage plots of HMGCS2 and PDK4 in MASH patients and HFD-fed mice were generated using SuperIntronic^30^.

### Statistical analysis

Statistical analysis was performed using R version 4.3.1. Student’s t-test was used for comparisons between two groups; Anova was used for comparisons between three or more groups, followed by post-hoc analysis with Tukey-HSD. All cell culture-based experiments have been conducted at least twice, and representative results were presented in Figures.

---

**Supplementary Figure 1.**
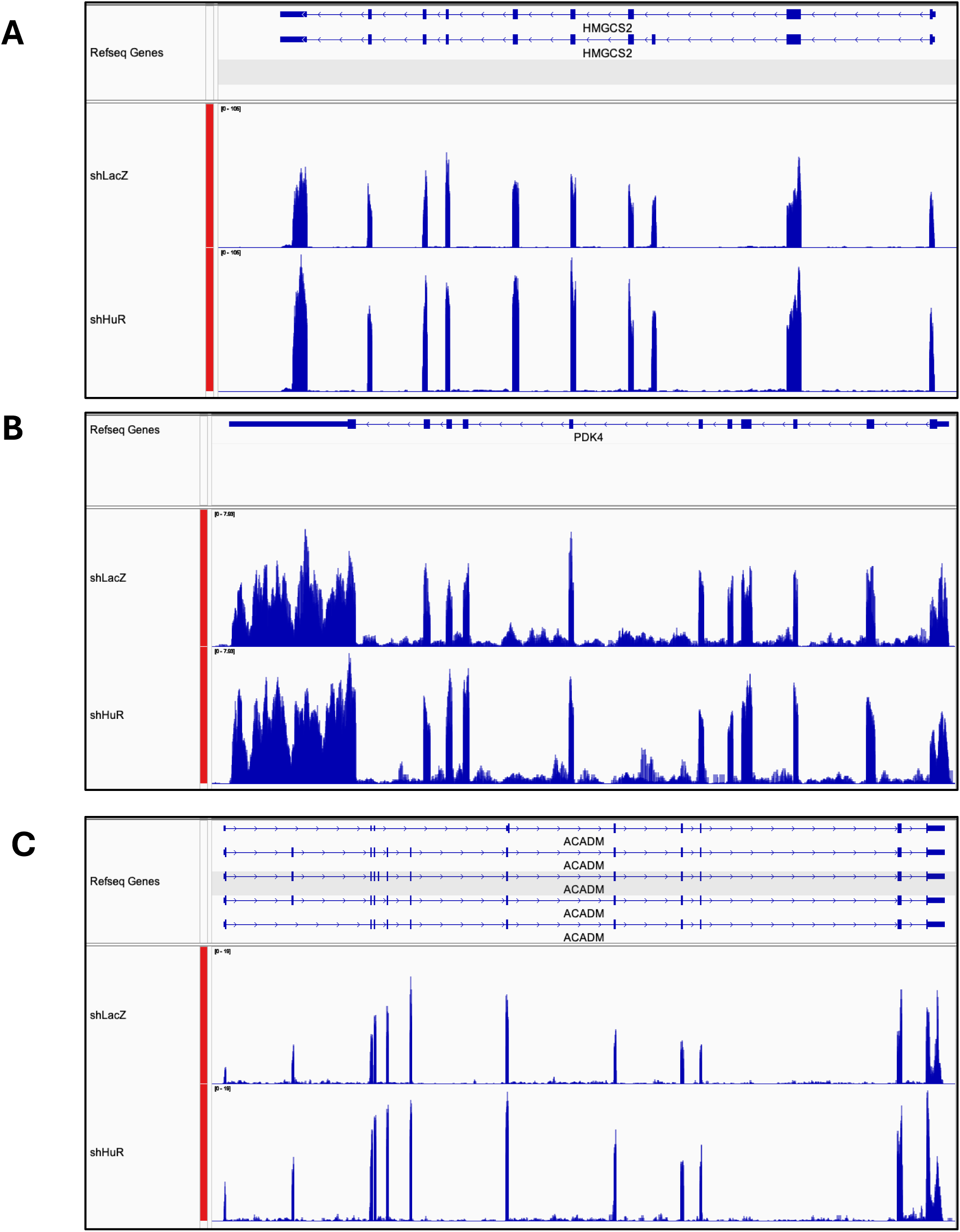
**A-C**, RNA-seq alignment for human HMGCS2, PDK4, and ACADM using data from humanized livers with control (shLacZ) or human HuR knocking down (shHuR) as presented by IGV genome browser. CPM-normalized bigwig files were loaded to IGV genome browser, and the signal of 4 samples of each group were overlayed. The data range was adjusted in the same range by using “Group Autoscale” function.

## Author Contributions

S.T. performed the experiments, bioinformatics analyses and analyzed the data. M.J. performed the experiments. S.T. and X.R. wrote the manuscript. X.R. conceived and supervised the study.

